# Reactive astrocytes - comprehending when neurons play 4’33”

**DOI:** 10.1101/189134

**Authors:** Vladislav Volman, Maxim Bazhenov

## Abstract

Homeostatic regulation is a powerful tool utilized by virtually all biological systems, brain included. Broadly speaking, each homeostatic process embodies two components: the sensing component, whereby a deviation is detected and quantified, and the effecting component that executes the homeostatic adjustment with the goal of alleviating the deviation. In the central nervous system, homeostatic plasticity has been suggested to play an important role in shaping the dynamics of single neurons and neuronal networks. However, existing “biophysical” models of homeostatic plasticity are exceedingly simplistic. These models usually describe the sensor component in terms of simple averaging over neuronal activity and offer no explanation of the relevant biochemical pathways. Here, we attempt to fill this gap in our understanding of homeostatic plasticity by proposing a biophysical framework to explain detection of prolonged synaptic inactivity that may occur in some scenarios of brain injury. We propose that sensing of, and response to, synaptic inactivity involves detection of the extracellular glutamate level and occurs via the activation of metabotropic glutamate receptors (mGluRs), while the inactivity-induced synthesis of one of the homeostatic plasticity effectors, tumor necrosis factor alpha (TNFα), serves as an effecting component of the system. This model can help to explain the experimental observations linking prolonged neuronal inactivity to TNFα signaling. Importantly, the proposed signaling scheme is not limited to mGluRs and astrocytes, but rather is potentially applicable to any cells expressing receptors that activate the relevant G protein units. The proposed signaling scheme is likely to be useful for developing pharmacological interventions targeting homeostatic plasticity pathways.

**Author summary:** Homeostatic regulation refers to the self-regulation ability of a system aimed at remaining in the same (or nearly the same) state. Homeostatic plasticity, a form of homeostatic regulation that arises in the context of neural dynamics, has been shown to shape neuronal and network dynamics by attempting to maintain physiological levels of neuronal activity in brain networks. Thus, it changes the excitation-inhibition balance in a network when sufficiently large and long-lasting deviations from physiological levels of activity are detected. However, the biophysical mechanisms of homeostatic plasticity and in particular those related to the ability to sense neuronal inactivity, remain elusive. We propose a feasible biophysical model of a homeostatic sensor, in which the sensing of inactivity depends on the presence and activation of G-protein-coupled receptors with different activation thresholds. In this model, the activation of the higher-threshold receptor suppresses the detection of inactivity, while the activation of the lower-threshold receptor promotes the detection of inactivity. The proposed model helps to explain a growing body of experimental data relating synaptic inactivity to production of homeostatic plasticity effectors such as tumor necrosis factor alpha. Although we consider a specific case study of metabotropic glutamate receptors on astrocytes, the model conclusions are likely to be applicable to any cells expressing receptors that activate the relevant G protein units.

## Introduction

The score of the magnum opus of John Cage, “4 minutes and 33 seconds”, instructs the performer not to play any instrument during that period, prompting the audience to comprehend the “sounds of silence”.

Extended periods of neuronal “silence” (inactivity) are a common feature of traumatic brain injury (TBI), arising as a result of neuronal, axonal, and synaptic injury [1,2]. During these post-traumatic “silence” periods, a number of physiological and anatomical changes may occur in the injured brain, aiming to offset abnormal neuronal activity incurred by traumatic events. Clinical data and *in vitro* models of TBI suggest that astrocytes, a subtype of glial cells, play a decisive role in the post-traumatic reorganization of neuronal circuitry [3]. Brain injury, induced either by mechanical or chemical means, triggers “reactive gliosis”, a complex multi-faceted process during which astrocytes undergo dramatic morphological and functional changes [3]. In response to injury, astrocytes synthesize and release plethora of molecules, which are likely to be critical in post-traumatic recovery but also in post-traumatic pathogenesis [3,4]. Notwithstanding the mounting evidence of astrocytic involvement in post-traumatic reorganization and pathogenesis, the biochemical pathways linking injury-induced neuronal inactivity to the synthesis and release of regulatory molecules by reactive astrocytes remain elusive.

One of the most consistent clinical findings pertaining to reactive gliosis is that of dramatically increased levels of tumor necrosis factor alpha (TNFα) a relatively short time following the traumatic event [5]. The timescale and extent of TNFα release vary and are likely to depend on the injury severity, with some injury models yielding high levels of extracellular TNFα as early as 1 h post-injury [6]. TNFα is an important mediator of neuroinflammation, and has been broadly associated with several neuropathological conditions. For example, high levels of TNFα were shown to correlate with cognitive impairment in a mouse model of multiple sclerosis [7]. In culture models, glial TNFα, synthesized in response to chronic neuronal inactivity, has been shown to scale glutamatergic and GABAergic synaptic conductance, thus altering network excitability [8,9]. Computational modeling studies further suggest that injury-induced synaptic scaling by glial TNFα might contribute to post-traumatic epileptogenesis (PTE) by breaching the excitation/inhibition balance [10,11]. Thus, understanding how injury-induced neuronal and synaptic inactivity leads to the glial synthesis and release of TNFα could further our understanding of the etiology of PTE.

Clinical models of TBI suggest that the post-traumatic increase in TNFα may be associated with a reduction in cyclic adenosine monophosphate (cAMP) [5]. This reasoning is supported by biochemical studies showing that suppression/enhancement of cAMP can up/down-regulate the synthesis of TNFα [5,12]. The down-regulating effect of cAMP on TNFα is mediated by cAMP-activated protein kinase A (PKA) suppression of nuclear factor kappa beta (NFkB), which controls the cellular synthesis of TNFα [13,14]. Thus, cAMP appears to be a critical causative factor in the astrocytic response to injury, but how does injury-induced inactivity induce cAMP variations remains largely unknown. Importantly, astrocytic cAMP is up-regulated by the activation of membrane receptors that couple to the G_q,s_ proteins, while cAMP accumulation is suppressed by the activation of receptors coupling to the G_i_ protein [15]. Astrocytes express a variety of G-protein-coupled receptors (GPCRs), among them adenosine receptors (ARs) and metabotropic glutamate receptors (mGluRs) [16,17], and extracellular application of either adenosine or glutamate (Glu) has been shown to result in the cAMP elevation via the activation of corresponding GPCRs [16,18]. This suggests that GPCR-mediated signaling is likely to account, at least partially, for cAMP changes post-injury. In relation to injury-induced prolonged synaptic inactivity mGluRs are particularly appealing because Glu is the main signaling molecule utilized by neurons (whereas the primary source of extracellular adenosine is glial cells [19]).

To relate extracellular Glu to the astrocytic response to prolonged synaptic inactivity, we constructed a computational model that incorporated known biochemical signal transduction pathways mediating the astrocytic response to this signaling molecule. We further used this model to investigate the properties of the astrocytic response to immediate sequelae of TBI, modeled as a chronic reduction in extracellular Glu in the proximity of astrocytic mGluRs. The model suggests that the presence of the two types of G protein pathways (the G_i_-protein-related cAMP suppression pathway and the G_q,s_-protein-related cAMP promotion pathway) and their activation by astrocytic mGluRs (and/or astrocytic ARs) can explain neuronal inactivity-induced variations in astrocytic cAMP. This, in turn, suggests that rather than targeting individual receptors as scapegoats, pharmacological models should consider common cellular circuits involved in pathogenesis.

## Results

Because excitatory synaptic transmission at central synapses is largely mediated by Glu, and because G-protein-coupled mGluRs are ubiquitously present on astrocytes, in this study we sought to determine the connection between extracellular Glu and astrocytic production of TNFα. To this end, we compiled several pieces of experimental evidence to build a biophysical computational model that accounted for sub-cellular pathways of the astrocytic response to changes in extracellular Glu. We assumed that: 1) In the healthy tissue, astrocytes express mGluR3 and mGluR5, with the mGluR3 activation affinity being higher (K_mGluR3_ = 0.5–10 μM) than the mGluR5 activation affinity (K_mGluR5_ = 2–14 μM) [17]; 2) The activation of mGluR5 promotes astrocytic calcium (Ca^2+^) via G_q,s_ signaling and production of inositol trisphosphate (IP_3_), whereas the activation of mGluR3 suppresses astrocytic cAMP via G_i_ signaling and suppression/reduction of the adenylyl cyclase (AC) activity [17,20]; 3) The Ca^2+^ and cAMP signaling pathways interact via the Ca^2+^-calmodulin (CaCAM) pathway [21]; 4) Low cAMP induces synthesis and release of TNFα through the suppression of the NFkB inhibitor ikB [5,13,14]. Schematic presentation of the modeled pathways is given in Fig 1 (see also Methods for details). The mechanisms downstream of cAMP involve transcriptional control, likely operate on timescales slower than those of second messenger dynamics, and are beyond the scope of the present study. Thus, we focused on understanding the putative link between neuronal inactivity and astrocytic cAMP levels.

**Fig 1.**
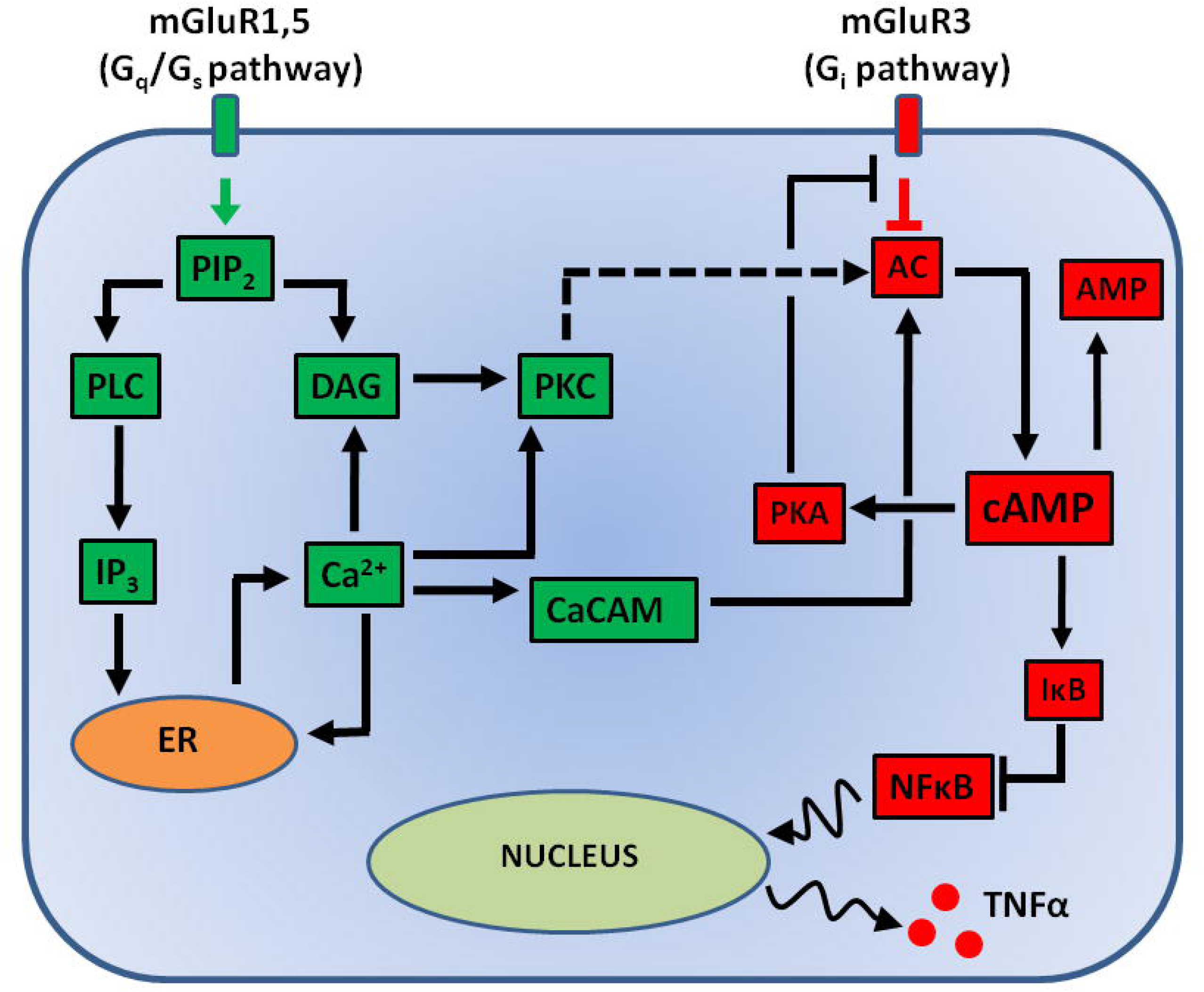
Schematic of modeled biochemical pathways linking synaptic inactivity to astrocytic TNFα.

### Steady state level of astrocytic cAMP depends on extracellular Glu

As a first step, we investigated the response of our model astrocyte to extracellular Glu. While *in vivo* extracellular levels of Glu likely fluctuate as a result of stochastic synaptic activity, constant extracellular Glu is a useful approximation that is often used in *in vitro* models. In the following, we assumed that mGluR3 (mGluR5) receptors have a relatively high (low) affinity to Glu [17]. Fig 2 shows that, for constant extracellular Glu, the temporal dynamics of astrocytic cAMP in the model strongly depended on the extracellular Glu concentration ([GLU]_EXT_). For low levels of [GLU]_EXT_ the concentration of cAMP ([cAMP]) remained constant, while higher [GLU]_EXT_ led to the appearance of [cAMP] oscillations (Fig 2A1,2) and higher time-averaged levels of [cAMP] (Fig 2B), suggesting the existence of a threshold (bifurcation) point associated with [cAMP] dynamics. This threshold dependence of [cAMP] on [GLU]_EXT_ reflected a similar relation between the astrocytic free Ca^2+^ concentration ([Ca^2+^]) and [GLU]_EXT_ (Fig. 2D1,2 and 2E), and was driven by the difference between the mGluR3 vs. mGluR5 affinities to Glu. Indeed, at low levels of [GLU]_EXT_, the cAMP response was mainly shaped by mGluR3, owing to the relatively high Glu affinity of these receptors, which suppressed AC and [cAMP] (Fig 1). In this regime, [Ca^2+^] and [cAMP] exhibited opposing trends with respect to [GLU]_EXT_. [Ca^2+^] increased, while [cAMP] decreased, with increasing [GLU]_EXT_. On the other hand, at higher levels of [GLU]_EXT_, the activation of mGluR5 positively affected [cAMP] via the CaCAM pathway, leading to the activation of AC and resulting in higher [cAMP], while the inhibitory effect of mGluR3 was saturated (Fig 1). This suggests that [GLU]_EXT_-dependent competition between the G_i_ (controlled by mGluR3) and G_q,s_ (controlled by mGluR5) pathways might determine astrocytic TNFα levels via [cAMP] regulation. Since our goal here is to predict under which conditions astrocytes can effectively detect neuronal inactivity, and since a strongly nonlinear dependence of [cAMP] on [GLU]_EXT_ (such as shown in Fig. 2B) would support this property of astrocytes, in the following we asked which properties of the G_i_ and G_q,s_ signaling pathways determine the sensitivity of the cAMP production mechanisms to extracellular Glu.

**Fig 2.**
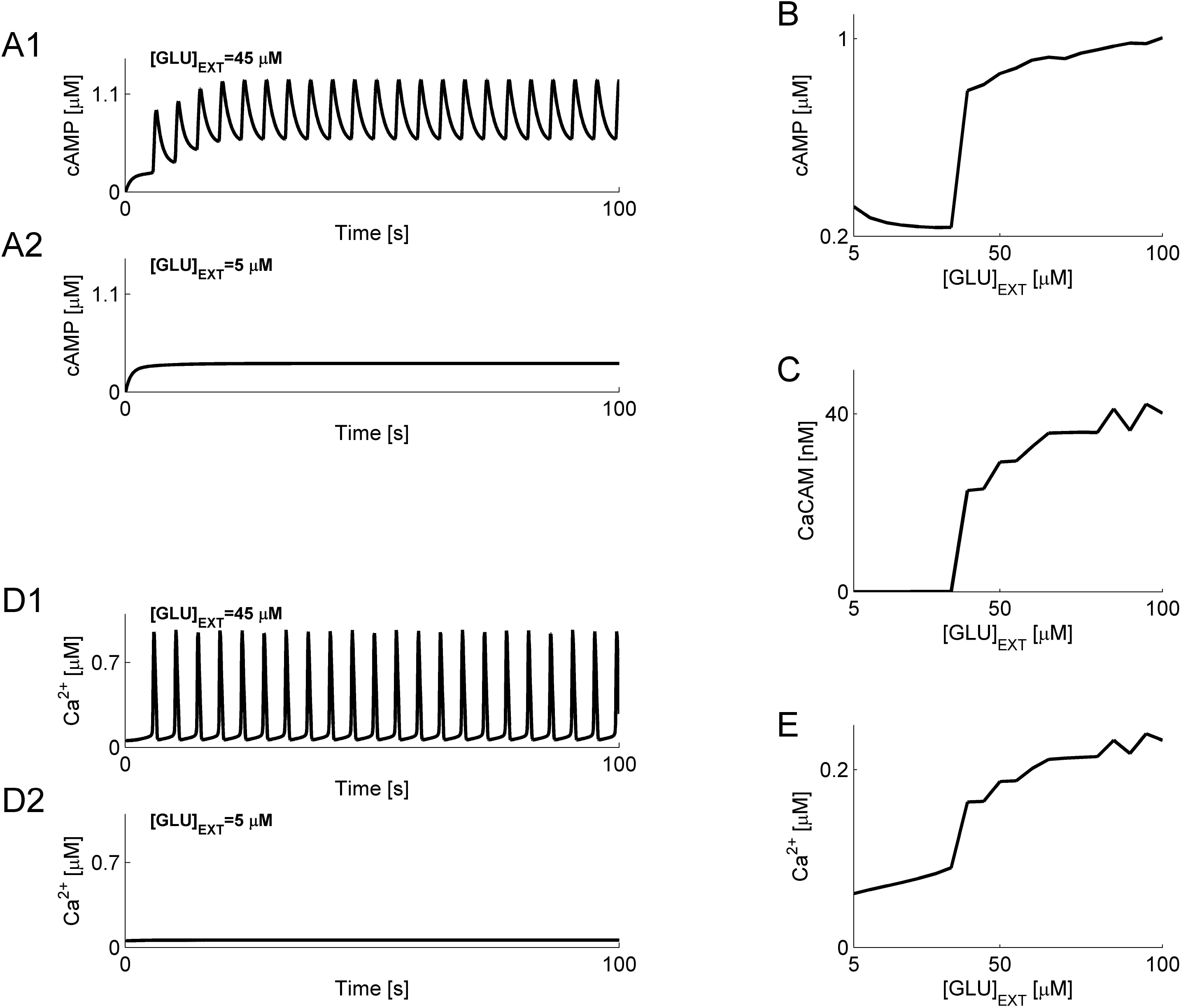
Dynamics of [Ca^2+^] and [cAMP] in the model astrocyte, under constant [GLU]_EXT_. (A) [cAMP] vs. time, for high (A1, [GLU]_EXT_ = 45 μM) and low (A2, [GLU]_EXT_ = 5 μM) [GLU]_EXT_. (B) Time-averaged [cAMP] vs. [GLU]_EXT_. (C) Time-averaged [CaCAM] vs. [GLU]_EXT_. (D) Free astrocytic [Ca^2+^] vs. time, for high (D1, [GLU]_EXT_ = 45 μM) and low (D2, [GLU]_EXT_ = 5 μM) [GLU]_EXT_. (E) Time-averaged free astrocytic [Ca^2+^] vs. [GLU]_EXT_.

### cAMP signaling pathway controls the discriminability of neuronal activity states

In addition to the modulation via the CaCAM pathway (related to the activation of mGluR5, see previous section), AC can also be modulated by the cAMP pathway. Indeed, activation of mGluR3 suppresses [cAMP], which implies lower levels of cAMP-activated protein kinase A (PKA). Because PKA downregulates the negative effect of mGluR3 activation on AC (Fig 1 and Equation (15)), activation of PKA (e.g., by cAMP) is expected to further facilitate cAMP production by weakening (through PKA) mGluR3 downregulation of AC (Fig 1). This feedback loop is also expected to contribute to shaping the dependence of [cAMP] on [GLU]_EXT_. In our model, the relative influence of this PKA-mediated AC modulation pathway depended on two parameters: the coupling strength, a_1_, between mGluR3 and AC, modulated by PKA (note that smaller a_1_ corresponds to stronger PKA-related decoupling of AC from mGluR3); and the half-activation concentration of PKA, p_T_, at which the feedback regulation was at the 50% of its maximal strength (with lower p_T_ yielding stronger PKA-related decoupling of AC from mGluR3). Thus, we next varied these two parameters to determine how they shape the relationship between [cAMP] and [GLU]_EXT_.

Fig 3A shows that the PKA-mediated modulation of the AC production was only effective for relatively low [GLU]_EXT_ (below ~40 μM). In this regime, the average [cAMP] was lower for higher a_1_ (corresponding to the weaker PKA decoupling of AC from mGluR3 and thus stronger mGluR3 suppression of AC) and the form of this dependence was significantly influenced by p_T_ (Fig. 3B1 and 3B2 for the low and high [GLU]_EXT_ regimes, respectively). To quantify this effect, we defined the discriminability of [cAMP] as D_I/T_ = 100·([cAMP]_H_-[cAMP]_L_)/[cAMP]_L_, where [cAMP]_L_ is the average [cAMP] in the low [GLU]_EXT_ regime ([GLU]_EXT_ in the 20–30 μM range), and [cAMP]_H_ is the average [cAMP] in the high [GLU]_EXT_ regime ([GLU]_EXT_ in the 80–90 μM range). With this definition, a higher discriminability implies a better astrocytic ability to translate [GLU]_EXT_ differences into distinct [cAMP] states and, therefore, a better ability to detect neuronal inactivity.

**Fig 3.**
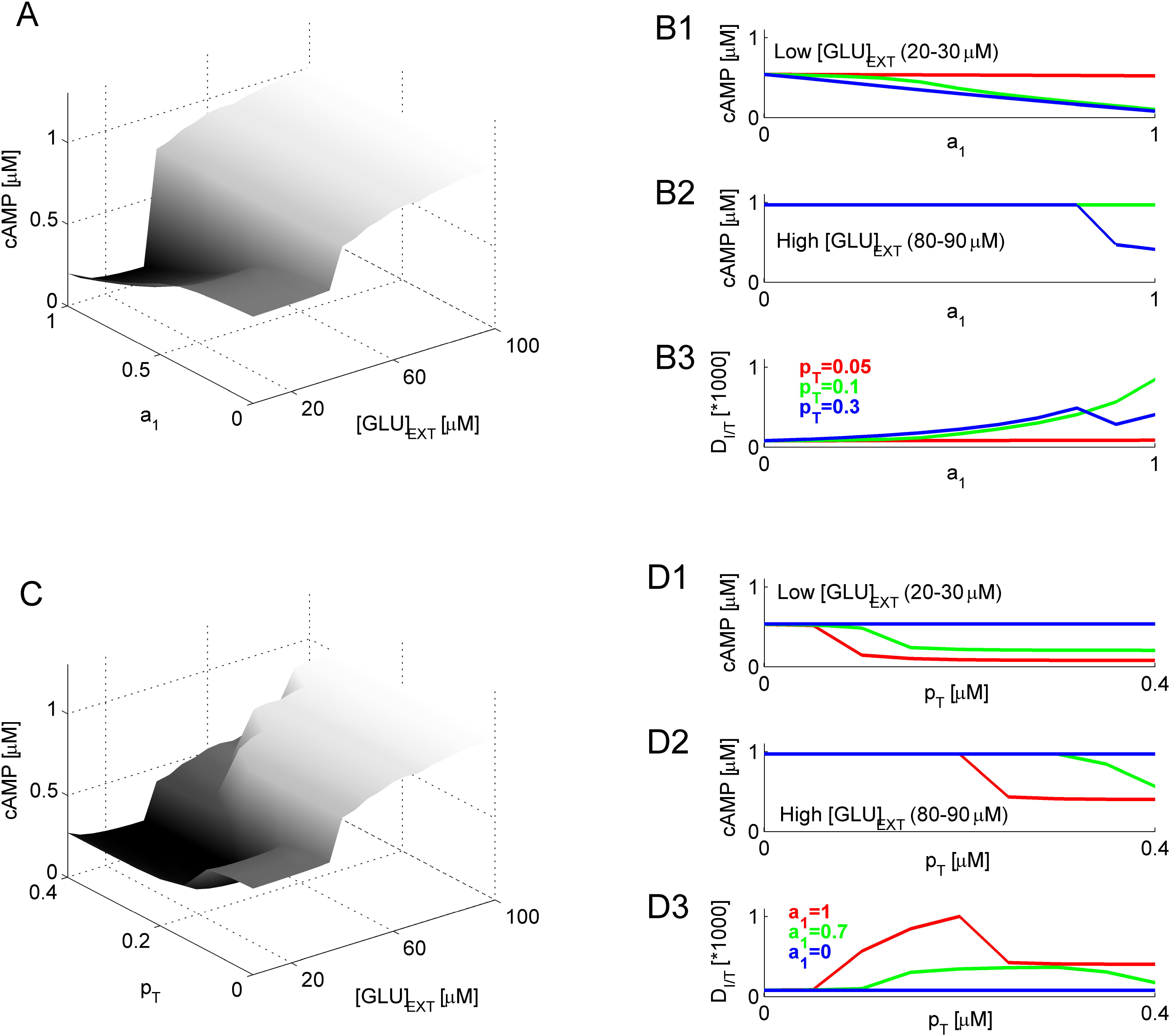
PKA regulation of mGluR3-AC coupling defines the discriminability of neuronal activity. (A) Time-averaged [cAMP] vs. the mGluR3-AC coupling strength, a_1_, and [GLU]_EXT_. (B) Time-averaged [cAMP] vs. a_1_, for different values of the half-activation parameter, p_T_, in low (B1, 20–30 µM) and high (B2, 80–90 µM) [GLU]_EXT_ regimes. (B3): Discriminability (defined in Text) vs. a_1_, for different values of p_T_. Data keys (shown in B3) are the same across all panels. (C) Time-averaged [cAMP] vs. p_T_ and [GLU]_EXT_. (D) Time-averaged [cAMP] vs. p_T_, for different values of a_1_, in low (D1, 20– 30 µM) and high (D2, 80–90 µM) [GLU]_EXT_ regimes. (D3): Discriminability (defined in Text) vs. p_T_, for different values of a_1_. Data keys (shown in D3) are the same across all panels.

Fig 3B3 shows that the discriminability increased with increasing the mGluR3-AC coupling strength, a_1_; this trend reflected a reduction in [cAMP] for higher values of a_1_. Interestingly, for sufficiently high a_1_ (strong mGluR3-AC coupling, or, alternatively, weak PKA downregulation of mGluR3-AC coupling) the discriminability was low for either very low or very high values of the half-activation parameter, p_T_, suggesting that there exists an optimal value of p_T_ (green curve in Fig. 3B3) for which the interaction is neither under- nor over- saturated. When plotted vs. p_T_ (for fixed values of a_1_), time-averaged [cAMP] exhibited the same qualitative dependence (Fig 3C and Fig. 3D1–3), confirming that the PKA signaling strongly affects [cAMP] in the low [GLU]_EXT_ regime, thus affecting the ability of the model astrocyte to discriminate between different levels of [GLU]_EXT_.

Our results so far suggest that astrocytic mGluR activation can strongly affect cAMP production in these cells. Thus, we next considered the contribution of the relative sensitivity to [GLU]_EXT_ (across the two receptor types, mGluR5 and mGluR3) to the astrocytic sensing of synaptic inactivity. Emerging evidence from the pharmacokinetic studies of mGluRs suggests that the sensitivity of these receptors to [GLU]_EXT_ can be modulated in a variety of ways (e.g., phosphorylation) [20]. Our simple model of mGluR activation neglected detailed binding/unbinding kinetics; in the model, receptor sensitivity to [GLU]_EXT_ was controlled by the receptor half-activation parameter (K_mGluR3_ and K_mGluR5_ for mGluR3 and mGluR5, respectively). A higher K_mGluR3/5_ corresponds to a lower receptor affinity to [GLU]_EXT_ (see Equations (1) and (15)). Thus, we varied the values of these parameters to determine their impact on the level of astrocytic [cAMP].

Fig 4A1 shows that varying K_mGluR3_ mostly affected the time-averaged levels of astrocytic [cAMP] in the low [GLU]_EXT_ regime ([GLU]_EXT_ < ~40 μM), but had a negligible effect on [cAMP] in the high [GLU]EXT regime ([GLU]EXT > ~40 μM). This is consistent with the idea that the G_i_ pathway controls the ability of the cAMP production mechanism to differentiate between different levels of neuronal activity (the discriminability was defined earlier as a relative change in [cAMP] across the low and high [GLU]_EXT_ regimes). Consistent with this, the discriminability decreased with increasing K_mGluR3_, and was higher for higher a_1_ (strong mGluR3-AC coupling, corresponding to weak PKA-related decoupling of AC from mGluR3) (Fig 4A2). On the other hand, the transition point from the low to high [GLU]_EXT_ regime (defined as the level of [GLU]_EXT_ at which the change in [cAMP] was the steepest) only weakly depended on K_mGluR3_ (Fig 4A3, note that the different curves practically coincide). Contrary to the effect of K_mGluR3_, variation in K_mGluR5_ strongly affected the transition point from the low to high [GLU]_EXT_ regime (Fig. 4B1 and 4B3), but exerted only a relatively weak effect on the discriminability of neuronal activity (Fig 4B2). Collectively, the results presented in Fig 4 suggest that different mGluRs may differently affect the ability of astrocytes to detect changes in neuronal activity, with Glu sensitivity of mGluR5 likely affecting the transition point between “inactive” and “active” neuronal states, and Glu sensitivity of mGluR3 likely defining the ability of astrocytes to discriminate between the above states by increasing or reducing the production of cAMP. This further suggests a critical role for the G_i_ pathway in determining the ability of astrocytes to detect neuronal inactivity.

**Fig 4.**
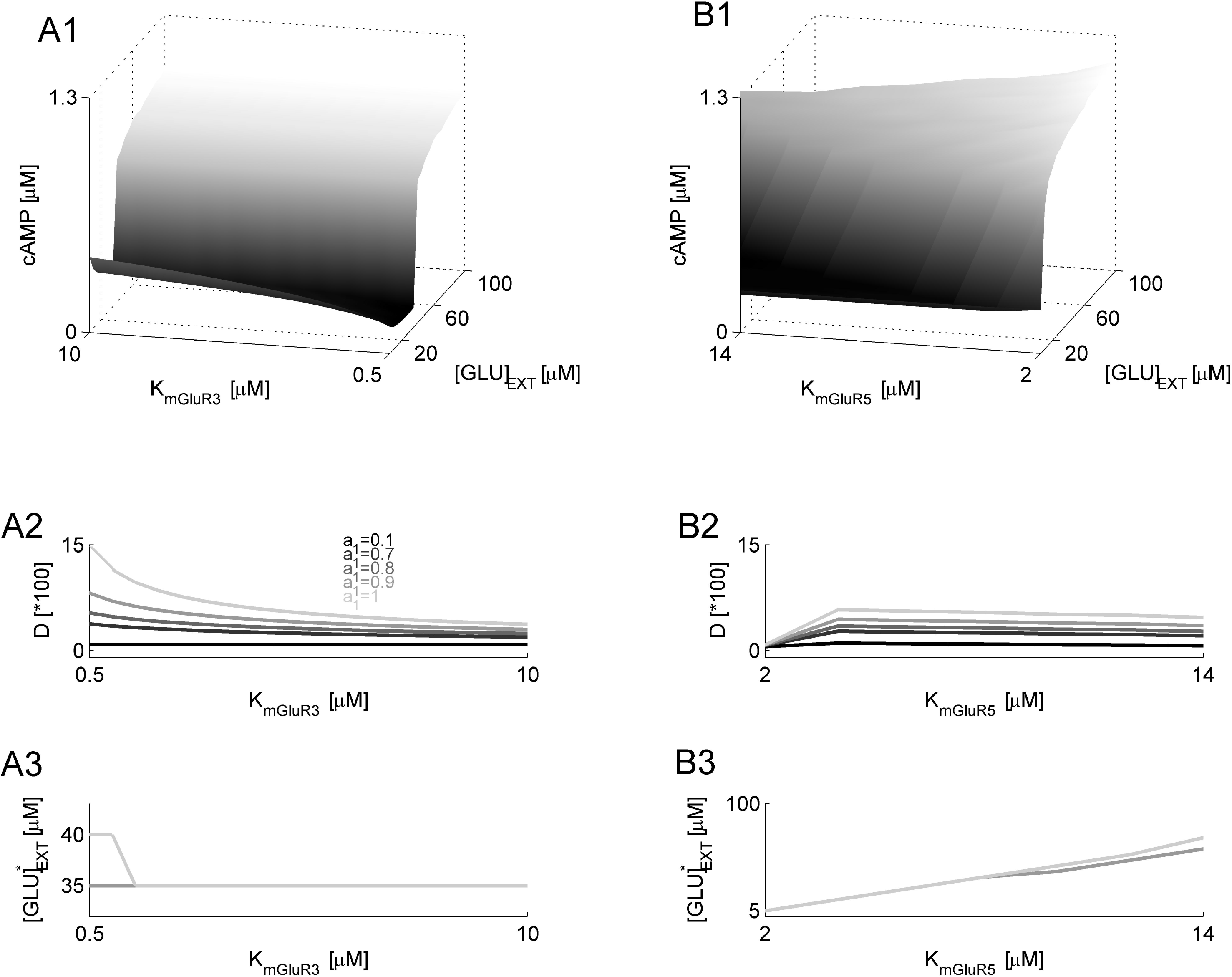
mGluR sensitivity to Glu defines astrocytic “eavesdropping” on neuronal activity. (A1) Time-averaged [cAMP] vs. the Glu half-activation of mGluR3, K_mGluR3_, and [GLU]_EXT_. (A2) Discriminability vs. K_mGluR3_, for different values of the mGluR3-AC coupling strength parameter, a_1_. (A3) “Low” to “high” [cAMP] transition point vs. K_mGluR3_, for different values of a_1_ (color code is the same as in A2). (B1) Time-averaged [cAMP] vs. the Glu half-activation of mGluR5, K_mGluR5_, and [GLU]_EXT_. (B2) Discriminability vs. K_mGluR5_, for different values of a_1_. (B3) “Low” to “high” [cAMP] transition point vs. K_mGluR5_, for different values of a_1_ (color code is the same as in B2).

### Modulation of astrocytic [cAMP] by synaptic activity-like fluctuations in [GLU]_EXT_

*In vivo*, the main contributor to [GLU]_EXT_ is synaptic activity. Because synaptic release of neurotransmitter from central synapses is highly stochastic [22], such stochasticity can translate into fluctuating [GLU]_EXT_, which might further affect the ability of astrocytes to eavesdrop on neuronal activity. Thus, we investigated the response of our model to fluctuating [GLU]_EXT_, a condition that mimics synaptic activity.

The extent of synaptic activity in our model was defined in terms of the rate of canonical synapse stimulation, ν_SYN_, and in terms of the increase, [GLU]_SYN_, in the extracellular Glu per synaptic event. Fig. 5A shows that the time-averaged [cAMP] in the model astrocyte positively correlates with both ν_SYN_ and [GLU]_SYN_. It appears, however, that synaptic stimulation rate (rather than [GLU]_SYN_) is a more critical determinant of [cAMP] oscillations (compare plots in Fig. 5A2–A5), because oscillations were absent in the sub-threshold stimulation regime (low ν_SYN_, Fig. 5A2, A4), regardless of [GLU]_SYN_. This dependence of astrocytic [cAMP] on the parameters of synaptic activity was qualitatively the same regardless of the presence or absence of the PKA regulation of mGluR3 (Fig 5, compare panels A, corresponding to a_1_ = 0.9, with panels B, corresponding to a_1_ = 0.1). However, for the stronger PKA-mediated decoupling of the G_i_ pathway (a_1_ = 0.1, Fig. 5B2–4), the level of [cAMP] in the regime of weak synaptic stimulation (low ν_SYN_) was significantly higher than that obtained for a similar stimulation in the a_1_ = 0.9 model (Fig. 5A2–4). This result again is consistent with our prediction that the G_i_ pathway primarily affects the discriminability of neural activity (i.e., discriminating between “weak” and “strong” synaptic activation regimes).

**Fig 5.**
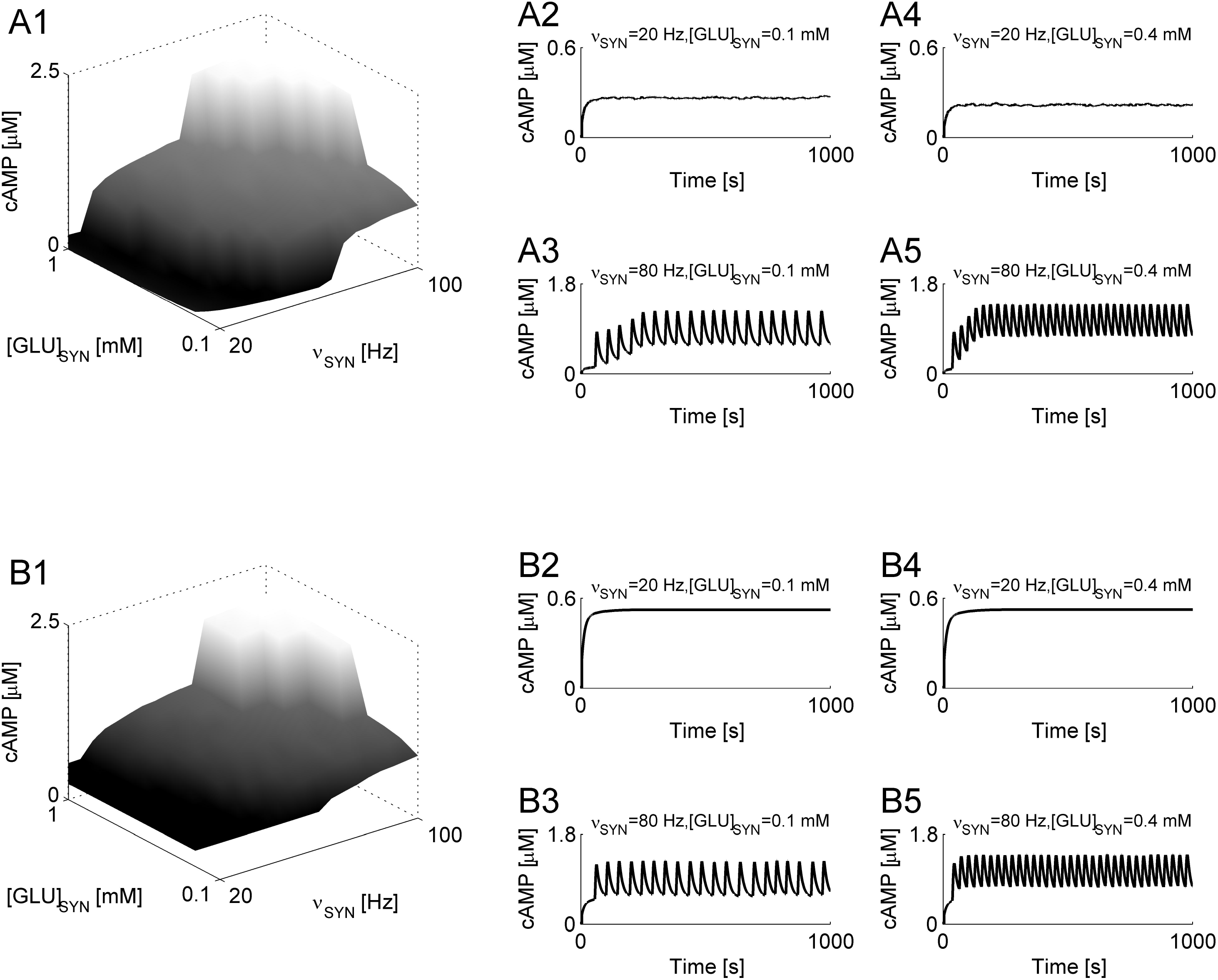
Synaptic-like fluctuations of [GLU]_EXT_ shape astrocytic [cAMP]. (A1) Time-averaged [cAMP] vs. the rate of synaptic activation, ν_SYN_, and the [GLU]_EXT_ increase per synaptic event, [GLU]_SYN_, for strong mGluR3-AC coupling (a_1_ = 0.9, corresponding to weak PKA-mediated decoupling). (A2)–(A5) Examples of [cAMP] temporal dynamics. (B1) Time-averaged [cAMP] vs. the rate of synaptic activation, ν_SYN_, and the Glu concentration increase per synaptic event, [GLU]_SYN_, for weak mGluR3-AC coupling (a_1 =_ 0.1, corresponding to strong PKA-mediated decoupling). (B2)–(B5) Examples of [cAMP] temporal dynamics.

## Discussion

On the sub-cellular level, Ca^2+^ and cAMP are engaged in an intricate interaction, often manifested as oscillations [23]. Whereas earlier studies mostly focused on the intracellular aspects of Ca^2+^–cAMP interaction, here we considered the activation of Ca^2+^ (G_q,s_, controlled by mGluR5) and cAMP (G_i_, controlled by mGluR3) pathways by extracellular agents, such as glutamate, and investigated the putative role of these pathways in determining the response of astrocytes to prolonged neuronal inactivity. Our model predicts that a reduction in [GLU]_EXT_ may trigger a dramatic reduction in astrocytic [cAMP] that may serve as a trigger for the commonly reported post-injury synthesis and release of TNFα. It further predicts critical roles for the relative contributions of the two Glu pathways (mGluR3- and mGluR5-activated) to the astrocytic ability to detect low levels of [GLU]_EXT_.

mGluR5 activation of the G_q,s_ transduction pathway increases IP_3_ and diacylglycerol (DAG) concentrations, which in turn increase free cellular [Ca^2+^] and activate protein kinase C (PKC), respectively [24]. In our model, the activation of mGluR5 affected [cAMP] via the CaCAM pathway (Methods); this choice was motivated by the existing biochemical data on calmodulin (CAM) activation that allowed us to reduce the number of unknowns in the model. Evidence suggests that in astrocytes, DAG-mediated activation of PKC can also significantly increase [cAMP] [25]. This interaction was not addressed in the present model, but because it is qualitatively equivalent to the studied one (according to which free intracellular [Ca^2+^] drives [cAMP]) we expect our conclusions (regarding the roles of different mGluRs in the [cAMP] regulation) to remain valid. IP_3_-induced Ca^2+^ release can also be facilitated by cAMP-dependent PKA [26], but by the same reasoning we expect our conclusions to remain valid after incorporating this mechanism.

In our model, AC was a critical mediator of the astrocytic response to prolonged neuronal inactivity, and we assumed that AC is inhibited by the activation of mGluR3 and modulated by Ca^2+^ via the CaCAM pathway. Nine isoforms of AC have been reported, all modulated by a variety of factors in different ways (reviewed in [27]). Of the nine known AC isoforms, AC1 and AC8 are the only ones that are likely to be both down-regulated by the activation of mGluR3 (via its G_iα_ unit) and up-regulated via the CaCAM pathway. At present, data on the expression of AC isoforms in astrocytes are not available. Our model predicts that either AC1 or AC8 are expressed in these cells and are linked to the astrocytic response to injury-induced prolonged neuronal inactivity.

The relative contributions of the two G-protein pathways (mGluR3- and mGluR5-activated) determined the production of cAMP in our model. At low levels of [GLU]_EXT_, AC was suppressed by the G_i_ pathway activation (controlled by mGluR3), which contributed to lowering the level of [cAMP]. With increasing [GLU]_EXT_, the G_q,s_ pathway (controlled by mGluR5) was activated, leading to a sharp increase in [cAMP]. Because the level of [cAMP] affects the synthesis and release of TNFα through the NFkB pathway (with high [cAMP] acting to suppress TNFα), our model proposes a link between low [GLU]_EXT_ (reflecting injury-induced prolonged synaptic inactivity) and synthesis and release of TNFα by astrocytes. We note that the abundance of astrocytes and their receptor makeup may vary across different brain regions/structures, making the proposed inactivity sensor more or less pronounced. This may be manifested as different propensities of brain regions/structures to mount TNFα response in response to the same level of injury. Whether or not this is the case can be determined by performing experiments to monitor TNFα while controlling for the severity of inactivity. In the meantime, although the quantitative specifics of glial response may be brain-region dependent, our model offers a generic causal link to explain the emergence of astrocytic neuroinflammatory response.

Chronic neuronal/synaptic inactivity that results in the astrocytic release of TNFα (as addressed by the present model) may be induced, for example, by a loss of functional afferent inputs that often occurs after TBI [28]. Evidence suggests that in the hippocampus TNFα affects synaptic neurotransmitter release [7,29] and can induce cognitive impairment. In general, TNFα appears to mediate or promote homeostatic synaptic scaling [8,9] and computational models [11,30,31] support the idea that such synaptic scaling can take part in PTE. Thus, our hypothesis provides a biochemical link between chronic neuronal inactivity and post-traumatic paroxysmal activity. Importantly, according to our hypothesis, the transition to paroxysms would occur as a result of low levels of [GLU]_EXT_. Mesial temporal lobe epilepsy (MTLE) was also postulated to be related to astrocytes, with epileptic activity arising owing to the excess of extracellular Glu [32]. This may appear to contradict our conclusions regarding the mechanisms of PTE. It is interesting, however, that the expression of both astrocytic mGluR3 and mGluR5 was found to be significantly up-regulated after seizures in an animal model of MTLE [33], as well as following kainate-induced seizures [34]. In our model, such up-regulation would correspond to a leftward shift of the [cAMP] vs. [GLU]_EXT_ curve (Fig 2), along with lower [cAMP] in the low [GLU]_EXT_ regime (increased discriminability); this change in turn would translate into higher astrocytic levels of [cAMP] (and thus lower levels of TNFα) in response to a lower [GLU]_EXT_. At the same time, astrocytic [cAMP] was found to be dramatically reduced after TBI [5]. It is thus tempting to speculate that astrocytic responses might be different for different epilepsies (caused by the excess/deficit of Glu, as in MTLE/PTE, respectively). Further, since TNFα inhibits the uptake of Glu by astrocytes [35], variations in astrocytic [cAMP] in response to chronic changes in synaptic activity might represent a homeostatic mechanism for maintaining physiologically reasonable levels of ambient Glu [36].

A role for cAMP in the attenuation of seizures was first proposed following the observation that induced seizures cause an elevation in cyclic nucleotide levels in brain [37]. Later studies confirmed that cAMP-elevating agents exert anti-inflammatory effects by enhancing the synthesis of anti-inflammatory cytokine interleukin 10 (IL-10) and concurrently suppressing the production of TNFα [12]. Finally, it was shown that [cAMP] is reduced following TBI [5]. Collectively, these studies suggest cAMP as a target for pharmacological control of astrocytic inflammatory response to brain injury. Our model further suggests that astrocytic [cAMP] can be modulated by the mGluR composition in these cells; thus, the model proposes a role for the mGluR activation in neuroinflammation, consistent with some recent proposals [38].

In our model, the astrocytic sensing of neuronal injury-induced inactivity was accomplished via the differential activation of astrocytic mGluRs. These receptors are coupled to G proteins, with mGluR5 coupled to the cAMP-promoting G_q,s_ and mGluR3 coupled to the cAMP-suppressing G_i_. Adenosine receptors (A_1_, A_2A_, A_2B_, and A_3_) were also suggested to be important in glial signaling [16]. Although all four types were detected in astrocytes, A_1_ (coupled to the G_i_ pathway, activation affinity of 0.31 μM [39]) and A_2B_ (coupled to the G_q,s_ pathway, activation affinity of 24 μM [39]) appear to be most prominently expressed. Basal extracellular adenosine levels are in the 25–250 nM range, and may increase following prolonged wakefulness [40] or localized brain injury [41] by up to 60-fold. Thus, prolonged wakefulness and/or localized brain injury are likely to suffice only to activate cAMP-suppressing (and TNFα-promoting) astrocytic A_1_ receptors, sharpening the effect described by our model. Note also that the link between prolonged wakefulness and TNFα synthesis, supported by the present model of G_i_-mediated astrocytic signaling, is consistent with the proposal of TNFα as a biomarker of sleep pressure [42].

## Methods

Our main motivation here was to delineate the putative biochemical mechanisms linking synaptic inactivity (low levels of extracellular Glu) to the TNFα response of astrocytes. Because mounting body of experimental evidence indicates that synthesis of TNFα is suppressed by elevated levels of cellular [cAMP] [5,12,13,14], we asked how [GLU]_EXT_ is linked to astrocytic [cAMP]. To address this question, we modeled the relation between the activation of different mGluRs (mGluR3 and mGluR5, known to be expressed in astrocytes) and astrocytic [cAMP]. An implicit assumption of our model was that synthesis of TNFα always increases with increasing [cAMP].

### Modeling the mGluR5 activation and the Ca^2+^ pathway

Activation of mGluR5 by extracellular Glu leads to the activation of phospholipase C beta (PLCβ) enzyme. PLCβ hydrolyzes phosphatidylinositol 4,5-bisphosphate (PIP_2_), resulting in the production of IP_3_ and DAG [24]. From there on, IP_3_ binds to its receptors on the endoplasmic reticulum (ER) and initiates the release of Ca^2+^ ions from ER to the cytosol, in the process termed Ca^2+^-induced Ca^2+^ release (CICR), while DAG activates PKC; however, the latter activation requires the presence of free Ca^2+^.

We assumed that the dependence of [PLCβ] on [GLU]_EXT_ (via the activation of mGluR5) in the proximity of receptors could be described by the following Hill function:

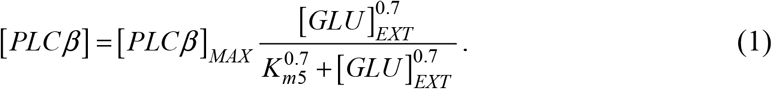

The kinetics of IP_3_ were

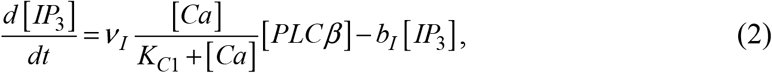

where **V**_I_ is the peak rate of IP3 production by PLCβ and free Ca^2+^, and bI is the rate of IP3 degradation. Detailed models of IP3 metabolism show that at physiological concentrations of IP_3_ degradation occurs via phosphorylation by the IP_3_ 3-kinase and can be effectively approximated as a linear process [43]. The model parameters were: **v**_I_ = 3.2 s^−1^, K_C1_ = 1 μM, b_I_ = 2 s^−1^. The peak [PLCβ] (obtained under the condition of Glu saturation of mGluR5) was set to [PLC(3]_MAX_ = 2.3 μM.

Ca^2+^ signaling in astrocytes is critically shaped by CICR; thus, we used the Li-Rinzel model [44], according to which the free intracellular [Ca^2+^] evolves as

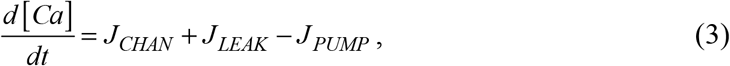

and the [Ca^2+^] fluxes are

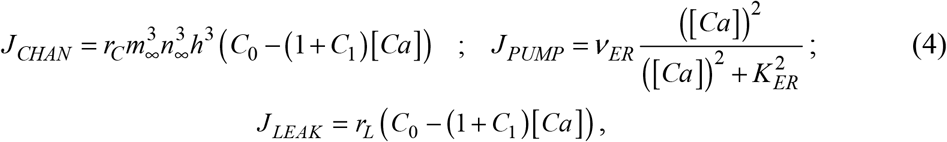

With

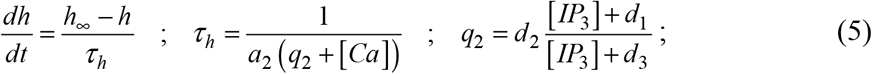

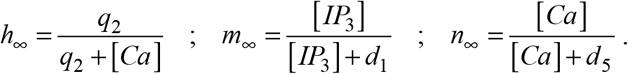

Our model of mGluR5-mediated Ca + signaling is clearly just an approximation of a complex web of regulatory loops and feedbacks. For example, cAMP-activated PKA can significantly affect intracellular [Ca^2+^] [23] and so can a regulatory system of IP3 metabolism [43]. Here, however, we were interested in the time-averaged long-term effect of [GLU]EXT on [cAMP]. Thus, since we considered the time-averaged values as a proxy to cellular responses, the exact and elaborate details of the signaling scheme became less important, allowing us to focus only on the core signaling pathways. The parameters of the model astrocytic [Ca^2+^] dynamics were: r_C_ = 6 s^−1^, V_ER_ = 0.9 μM-s^−1^, K_ER_ = 0.1 μM, r_L_ = 0.11 s^−1^, d_1_ = 0.13 μM, d_2_ = 1.049 μM, d_3_ = 0.9434 μM, d_5_ = 0.08234 μM, a_2_ = 0.2 s^-1^, C_0_ = 2 μM, C_1_ = 0.185 μM.

Several lines of evidence (reviewed in [27]) suggest that the function of AC (and consequently the levels of [cAMP]) is modulated by [Ca^2+^] via the activation of CAM complex. Ca + binds CAM in four steps, and the corresponding kinetic reactions are

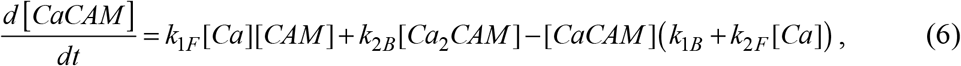

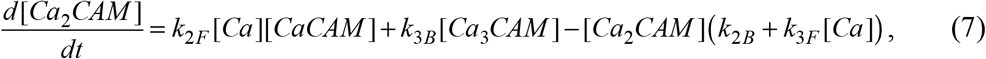

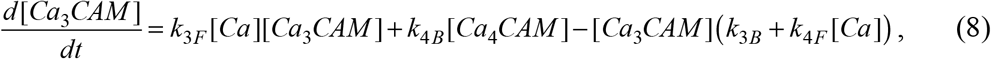

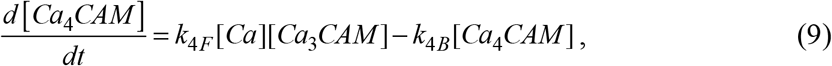

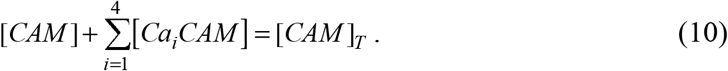

The binding/unbinding rates of Ca^2+^ to CAM are much faster than the timescales we consider (the period of a typical [Ca^2+^] oscillation is ~10s of seconds). Thus, we used a pseudo-steady state approximation to rewrite the levels of different binding steps as

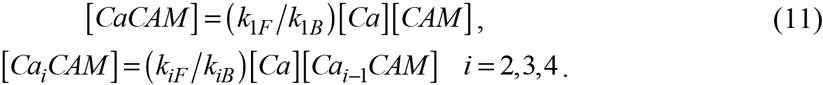

Earlier modeling and physiology studies [21,45] suggested that the third ([Ca_3_CAM]) and the fourth ([Ca_4_CAM]) steps of the CaCAM pathway are exclusively and equally effective in mediating the effect of [Ca^2+^] on [cAMP]. We followed the conclusions of these earlier studies by assuming that only the third and the fourth steps of the CaCAM pathway are important in the [cAMP] modulation, and that their contribution is equal. The kinetic parameters of the different binding/unbinding steps were: k_1F_ = k_2F_ = 2.3 nM^-1^·s^-1^, k_3F_ = k_4F_ = 160 nM^-1^·s^-1^, k_1B_ = k_2B_ = 2400 s^-1^, k_3B_ = k_4B_ = 405000 s^-1^. The overall concentration of CAM was [CAM] = 10 μM.

### Modeling the cAMP pathway

The level of [cAMP] is determined by AC (that catalyzes cAMP production from ATP), and by phosphodiesterases (PDEs) that catalyze the decomposition of cAMP into adenosine monophosphate (AMP). The concentration of AC can be further affected by various factors (reviewed in [27]). We focused here on the possible role of mGluR activation in modulating the levels of [cAMP]. Free intracellular [Ca^2+^], which in our model was elevated following the mGluR5 activation and release of Ca^2+^ from intracellular stores, up-regulates AC via the CaCAM pathway [27]. On the other hand, mGluR3 downregulate AC [20]. In our model, these different effects of the mGluR5 and mGluR3 activation on [cAMP] were captured in the following equation that governed the dynamics of [cAMP]:

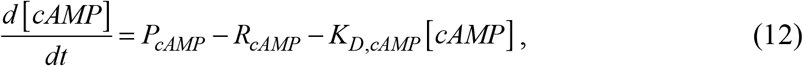

where P_cAMP_ describes the production of cAMP, R_cAMP_ describes the reduction in [cAMP] (due to the action of PDE), and K_D,cAMP_ models the rate of PDE-independent decay of cAMP. Each one of these terms is described in the following subsections.

cAMP activates PKA by binding to it. The dynamics of activated PKA (PKA*) concentration were described by the following equation, in which we implicitly assumed abundance of inactive PKA (all effectively lumped in the rate parameter, ν_a_^(1)^):

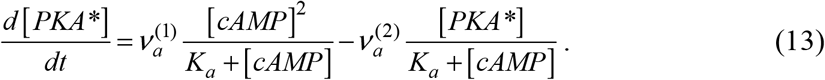

The parameters were [46]: v_a_^(1)^ = 810 ^3^ s ^1^, v_a_^(2)^ = 10 ^2^ μM-s ^1^, K_a_ = 10 ^-2^ μM.

### cAMP up-regulation

[cAMP] is up-regulated by AC, which in itself is driven by the activation of beta-adrenergic receptors. In turn, AC is modulated by [Ca^2+^] via the CaCAM pathway [27], and we followed Fridlyand et al. [45] in modeling this effect:

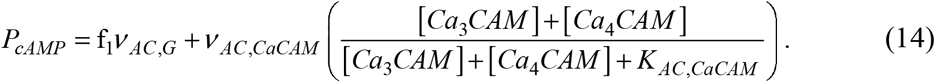

Activation of mGluR3 down-regulates AC [20]. Further, experimental evidence suggests that AC is likely to be regulated by the PKA phosphorylation of mGluR3, which decouples these receptors from their binding proteins [47]. We assumed that the feedback regulation of AC by the activated PKA could be described by the nonlinear function f_2_ = (1+exp(([PKA*]-p_T_)/0.4))^-1^, where p_T_ defines the threshold concentration of PKA for which the regulation is at the 50% of its maximal effect. We incorporated the available empirical data into our model by assuming that the dependence of AC on [GLU]_EXT_ in the mGluR3 proximity can be described by the following Hill function:

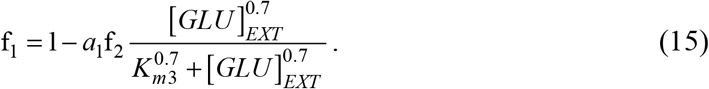

Kinetic parameters of [GLU]_EXT_ activation of mGluR3 are unknown at present; thus, we assumed the same exponent (0.7) as the one used for modeling the mGluR5 activation. To determine the robustness of our conclusions, we simulated the model with different values of mGluR3 activation exponent (in the 1–2 range), and obtained qualitatively the same results (simulation results not shown). The parameter a_1_ (in the 0–1 range) defined the strength of AC inhibition by the mGluR3 activation (the strength of mGluR3-AC coupling), and, as we show in Results, this parameter critically shaped the response of the model astrocyte to synaptic inactivity.

### cAMP down-regulation

[cAMP] down-regulation is primarily mediated by PDEs. Importantly, PDEs can be modulated via the CaCAM pathway as well. We followed Fridlyand et al. [45] in modeling the effect of PDEs on astrocytic [cAMP]. Thus, the term R_cAMP_, capturing the degradation of cAMP, was:

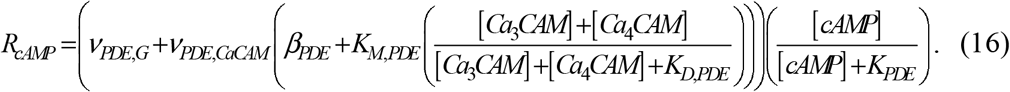

The parameters describing the production and decay of cAMP were [45]: V_AC,G_ = 0.048 μM-s^−1^, V_AC,CaCAM_ = 0.255 μM’s^1^, K_AC,CaCAM_ = 0.348 μM, V_PDE,G_ = 0.24 μM-s^−1^, V_PDE,CaCAM_ = 0.12 μM-s^−1^, β_PPDE_ = 0.4, K_M,PDE_ = 1.2, K_D,PDE_ = 0.348 μM, K_PDE_ = 3 μM, K_D,cAMP_ = 0.01 s^−1^.

### Extracellular Glu

We considered two different scenarios of [GLU]ext in investigating astrocytic response to injury-incurred prolonged synaptic inactivity. In the first scenario (which we term “the static scenario”) we investigated the response of the model astrocyte to constant (over time and space) concentrations of [GLU]ext. Conceptually, constant [GLU]ext can be thought of as resulting from temporal averaging over concentration fluctuations induced by synaptic events. Approximating [GLU]ext by a constant allowed us to conduct parametric space exploration, with constant [GLU]ext as the only measure of synaptic activity.

In a separate set of simulations, we considered temporally fluctuating [GLU]ext, mimicking synaptic activity. The Glu concentration in the mGluR proximity was:

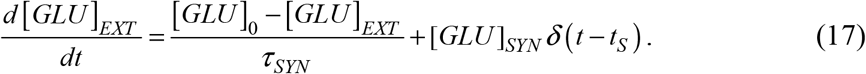

In Equation (17), synaptic events were generated at times *t_S_* according to a Poisson process, with the process rate of *ν_SYN_*. Each synaptic event incremented [GLU]_EXT_ in the mGluR proximity by [GLU]_SYN_, reflecting the number of Glu molecules that diffused to glial receptors following a synaptic release event. Vesicular concentration of Glu has been estimated to be 100 mM [36]. We assumed that up to 10% of synaptically released Glu diffuses to glial receptors; thus, the upper limit on [GLU]_SYN_ in the model was 10 mM. The characteristic time of [GLU]_EXT_ decay was τ_SYN_ = 10 ms. The quantity [GLU]_0 =_ 50 nM was the baseline [GLU]_EXT_ that would be obtained in the complete absence of synaptic activity.

